## Supplementary Figures for "A prefrontal cortex-lateral hypothalamus circuit controls stress-driven increased food intake"

### Supplementary Information

### Supplementary Figure Legends

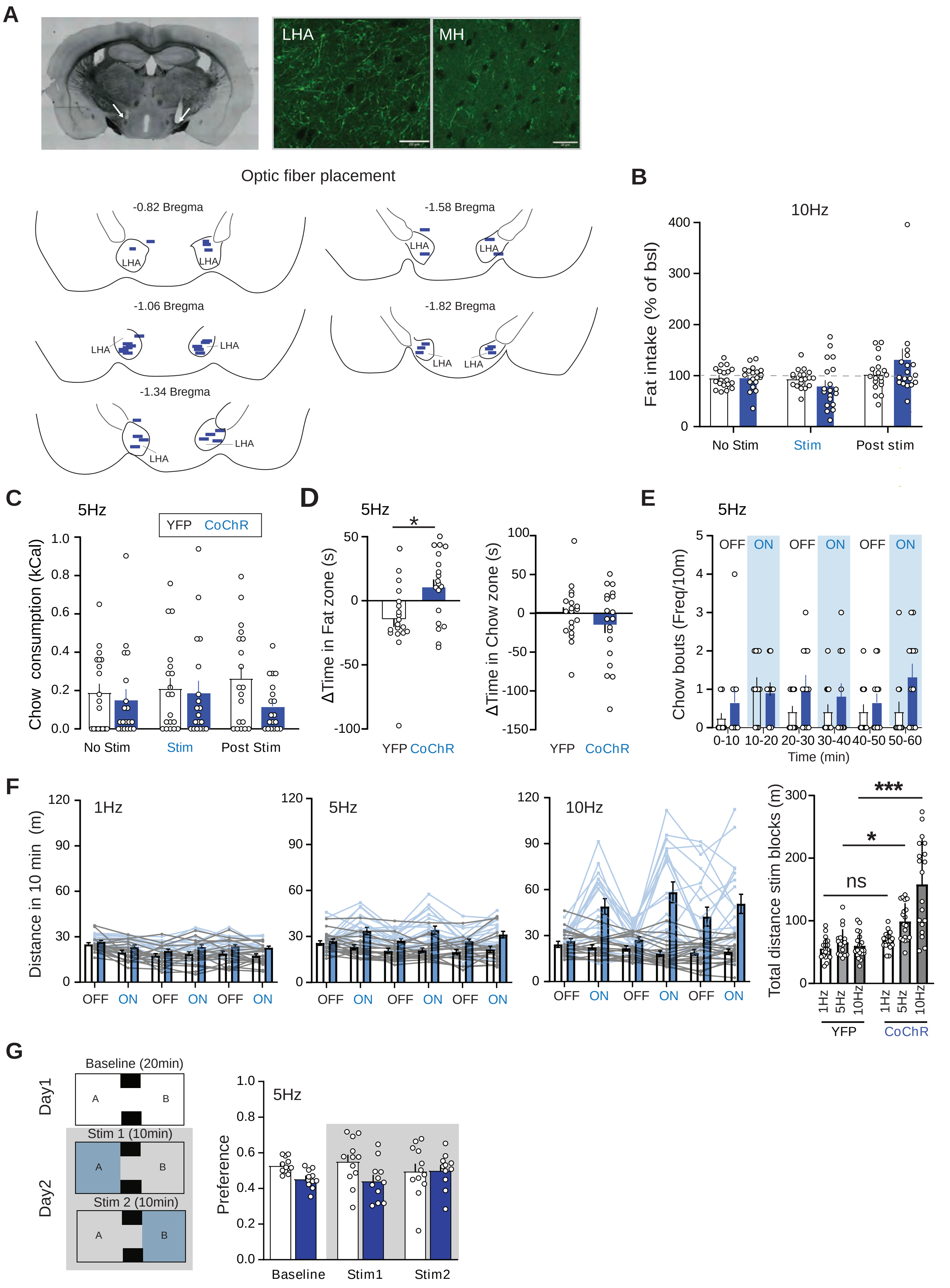


Supplementary Figure 1

**Supplementary Figure 1. Frequency dependent modulation of palatable food intake by mPFC-LHA pathway stimulation.** (**A**) All mice were histologically assessed for viral expression in mPFC, axonal fibers in LHA and correct placement of optic fibers. Optic fiber locations of all mice are drawn on the mouse brain atlas schematics. Displayed are all CoChR mice for this experiment (n=19). **(B)** Bar graph quantifying fat consumption normalized to intake of the first three baseline days for continuous 10 Hz stimulation. This did not affect food intake (YFP n=19, CoChR n=19, Two-way RM ANOVA, Day-Virus interaction, $F(2,72)=2.52$, $p=0.10$). (**C**) Bar graph quantifying 5Hz stimulation of mPFC-LHA pathway on chow intake (YFP n=19, CoChR n=19, Two-way RM ANOVA, Interaction Time-Virus, $F(2,72)=2.66$, $p=0.09$). (**D**) Bar graph showing effect of mPFC-LHA 5 Hz stimulation on time spent near the fat (YFP n=19, CoChR n=19, Mann Whitney test, $p=0.01$), and near chow (YFP n=19, CoChR n=19, unpaired t-test, $p=0.3$). (**E**) Bar graph quantifying the number of chow feeding bouts after mPFC-LHA stimulation at 5Hz (YFP n=12, CoChR n=12, Two-way RM ANOVA, Interaction time x virus, $F(5,110)=1.19$, $p=0.32$) (**F**) Bar graphs on effect of mPFC-LHA on locomotor activity displayed for laser ON and OFF block for the different frequencies. An increase in distance travelled during stimulation blocks was found with increasing frequencies (YFP n=12, CoChR n=12, Two-way ANOVA, Interaction Frequency-Virus, $F(2,108)=14.97$, $p<0.001$, Sidák's multiple comparison test YFP vs CoChR, 1Hz, $p=0.54$, 5Hz $p=0.01$, 10Hz $p<0.001$). **(G)** mPFC-LHA stimulation effect in the real-time place preference task. Preference is plotted as time spent in stimulated compartment as a fraction of total time spent in the stimulated and non-stimulated compartment (YFP n=12, CoChR n=11, 5Hz: Two-way RM ANOVA, $F(2,42)=2.32$, $p=0.11$). *= $p<0.05$; ***= $p<0.001$.

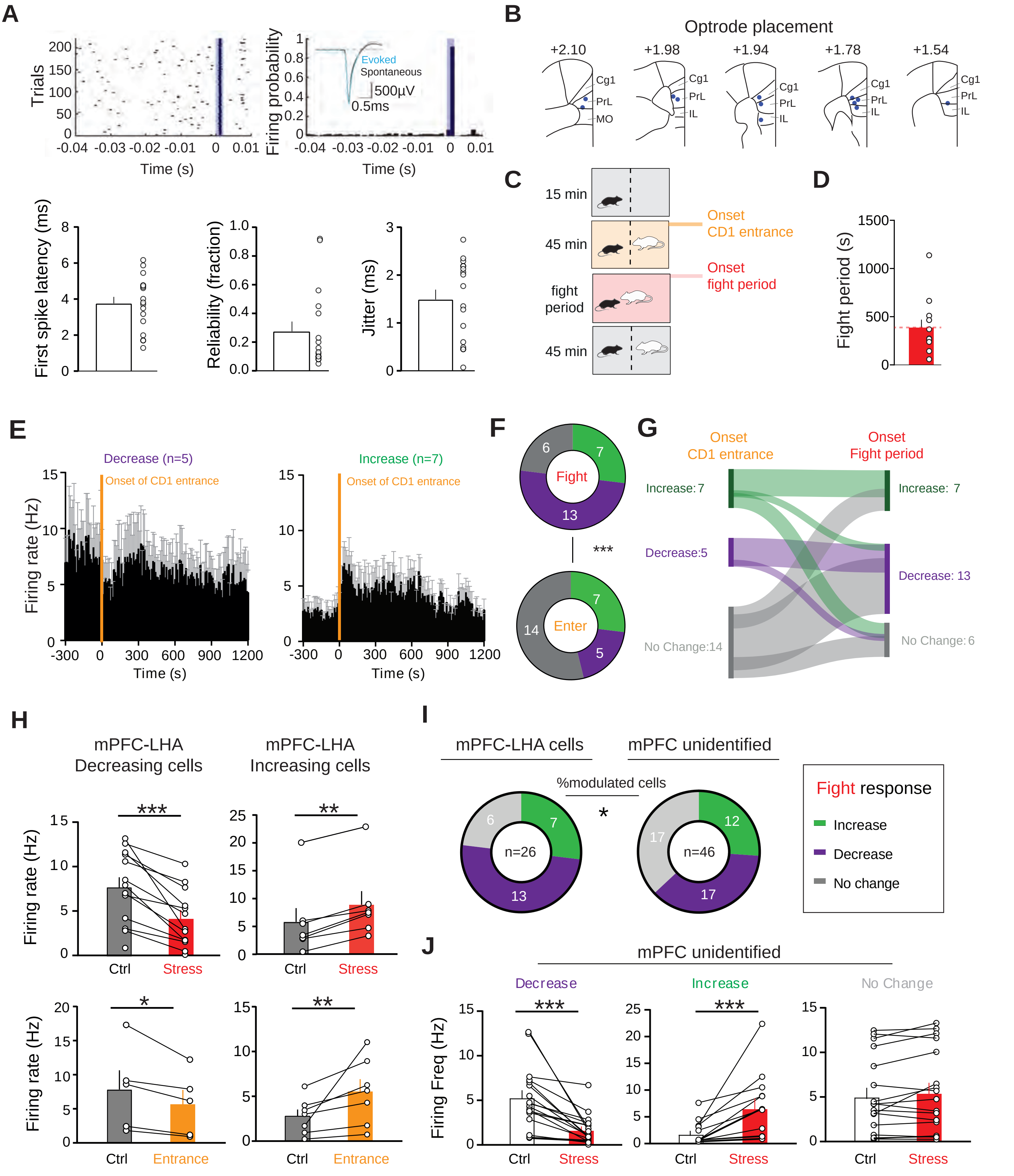


Supplementary Figure 2

**Supplementary Figure 2. mPFC-LHA neuronal activity is acutely modulated by social stress.** (**A**) Details and identification criteria for optogenetic identification of mPFC-LHA neurons. mPFC-LHA neurons were identified based on reliable, short-latency firing with low jitter after applying 2ms laser pulses for ≥200 trials. (**B**) Histological verification of optrode tip placement within mPFC (n=12 mice). (**C**) Schematic of different phases of exposure to social stress: recording alignment occurred to the onset of the physical interaction period or the timepoint when the CD1 was placed next to the test mouse separated by a barrier. **(D)** Bar graph with average exposure time of mice leading up to 20s of cumulative fight (n=12 mice). (**E**) Histogram showing the average response of those neurons which significantly decreased (n=5 neuron in 3 mice), or of those neurons that significantly increased (n=7 neurons in 5 mice) their firing after the CD1 mouse entered the cage (10s bins comparing 180 seconds right before and after onset of CD1 entrance). **(F)** Pie charts comparing within neuron response types for onset of fight period and CD1 entrance (n=26 cells in 12 mice, Chi-square test, Z=15.59, p=0.0004). **(G)** Sankey graph displaying the relation between the response types for the two different social stress situations of CD1 entrance without fight, and during the fight period (n=26 cells in 12 mice). **(H)** Bar graphs quantifying the average firing rate pre- and post CD1 entrance and onset of fight period for significantly modulated mPFC-LHA neurons grouped per response type (Fight period: decrease (n=13 cells in 9 mice), increase (n=7 cells in 7 mice); CD1 entrance: decrease (n=5 cells, 3 mice), increase (n=7 cells, 5 mice). **(I)** Comparison of response distributions of opto-tagged mPFC-LHA cells and non-identified mPFC (putative pyramidal) cells. Modulation (up or down) by social stress, binomial test, p=0.034. **(J)** Bar graphs quantifying the average firing rate pre- and post the onset of fight period for significantly modulated mPFC neurons grouped per response type, decrease (n=17 cells in 7 mice), increase (n=12 cells in 6 mice) and no Change (n=17 cells in 7 mice). *p<0.05, **p<0.01, ***p<0.001.

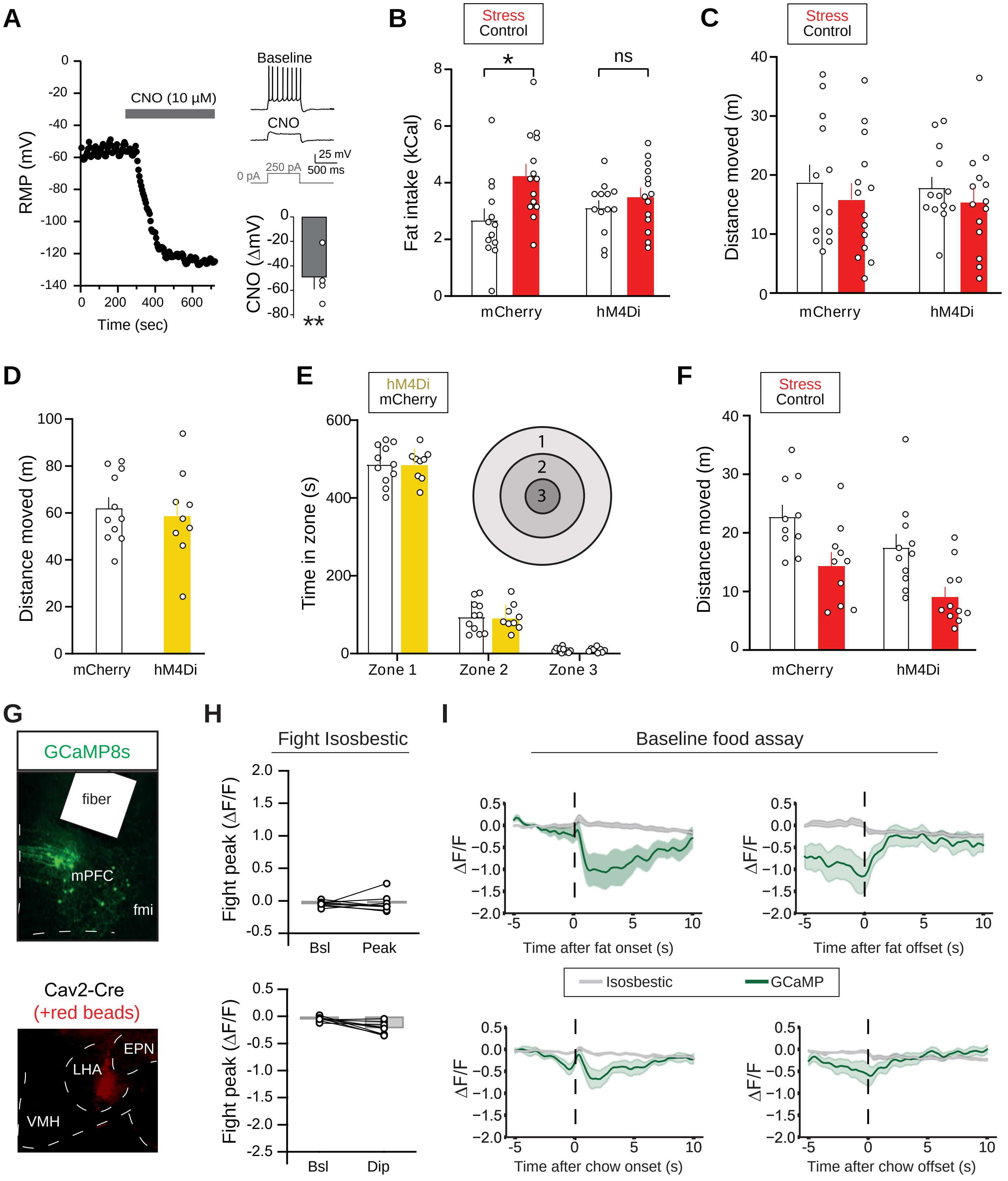


Supplementary Figure 3

**Supplementary Figure 3. Chemogenetic manipulation and fiber photometric recordings of mPFC-LHA pathway. (A)** Current clamp recordings in brain slices from mPFC-LHA neurons intersectionally virally targeted to express chemogenetic actuator hM4Di (see Fig. 3A). Left: Example mPFC-LHA cell where the resting membrane potential (RMP) hyperpolarized upon bath application of CNO. Top right: Example of the decrease in excitability of this cell in response to the same depolarizing current step during baseline and during CNO application. Bottom right: Bar graph quantifying the extent of hyperpolarization induced by CNO. **(B)** Absolute fat intake of mice during post-stress feeding assay. Mice ate more fat under stress conditions without inhibition of the mPFC-LHA pathway which was abolished by chemogenetic inhibition (mCherry Control n=13, hM4Di control n=13, mCherry Stress n=14, hM4DGi Stress n=14, 2 way ANOVA, main effect of virus, $F(1,50) = 8.12$, $p=0.006$, Sidák's multiple comparison test, mCherry control vs mCherry stress, $p=0.01$). **(C)** No difference in total distance moved in the food context under inhibition of the mPFC-LHA pathway after the 2 day social stress context (mCherry Control n=13, hM4Di control n=13, mCherry Stress n=14, hM4Di Stress n=14, Two-way ANOVA, main effect virus, $F(1,50) = 1.1$, $=0.30$). **(D)** mPFC-LHA inhibition did not affect total locomotor activity in an open field test (mCherry n=11, hM4Di n=9, unpaired t-test, $p=0.68$). **(E)** No differences were observed in anxiety measures of the open field. Mice spent an equal amount of time in all zones regardless of mPFC-LHA inhibition (mCherry n=11, hM4Di n=9, Two-way ANOVA, interaction Zone x Virus, $F(2,54) = 0.003$, $p=0.99$). **(F)** No interaction was found between mPFC-LHA inhibition and the effect of stress on locomotor activity in the food context after acute stress (mCherry Control n=10, hM4Di control n=11, mCherry Stress n=10, hM4Di Stress n=11, Two-way ANOVA, interaction Treatment x Virus, $F(1,38)=0.0003$, $p=0.99$). **(G)** Histological validation of fiber photometric recordings from the mPFC-LHA pathway. **(H)** Isosbestic controls for fight graphs (Fig. 3G). **(I)** Time courses for fat and chow interactions (onsets and offsets) prior to stress (or control) experiences. *$p<0.05$, **$p<0.01$, ***$p<0.001$.

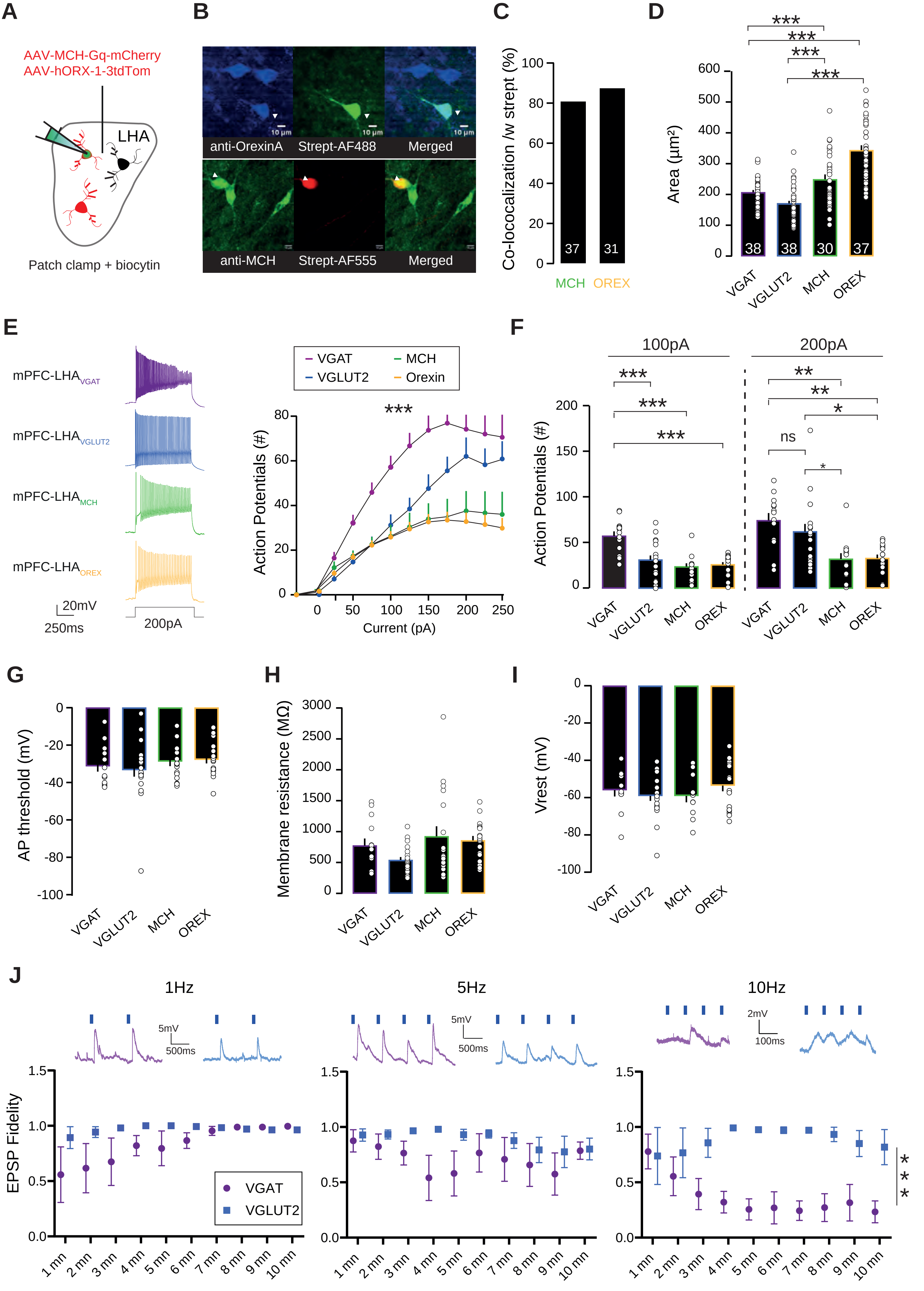


Supplementary Figure 4

**Supplementary Figure 4. mPFC neurons target multiple LHA subpopulations. (A)** Schematic of patch clamp experiments to validate targeting of $LHA_{OREX}$ and $LHA_{MCH}$ neurons. **(B)** Representative image of Orexin (top) and MCH (bottom) cells with viral and antibody cross-validation. Colocalization indicated with white arrows. Scale bar 10 µm. **(C)** Percentage of streptavidin-cell colocalized with antibodies (n MCH cells=37, 19 mice; n Orexin cells= 31, 10 mice). **(D)** Bar graph for soma area also shows an increase of size for MCH and Orexin neurons (n VGAT cells= 38, 2 mice, n VGLUT2 cells= 38, 2 mice, n MCH cells=30, 2 mice and n Orexin cells=37, 2 mice, One-Way ANOVA, $F(3,139)=37.739$, $p <0.001$). **(E)** Left: Example traces of action potential cells in response to a depolarizing current of 200pA in different LHA cell types. Right: Current-action potential relationships for the four distinct LHA populations (n VGAT cells=13, 7 mice, n VGLUT2 cells=20, 8 mice, n MCH cells=18, 9 mice and n Orexin cells= 18, 5 mice; Two-way RM ANOVA, Main effect of group, $F(3,65)=12.002$, $p<0.001$). (**F)** Bar graph for action potential number elicited by 100pA (left) and 200pA (right) current injection from the four LHA neuronal subsets (n as in E, Two-way RM ANOVA, Main effect of group at 100pA, $F(3,65)=11.343$, $p<0.001$ and Main effect of group at 200pA, $F(3,65)=8.355$, $p<0.001$). **(G)** Bar graph for the action potential threshold. **(H)** As (G), but for the membrane resistance. **(I)** As (G), but for the resting membrane potential across the 4 LHA populations. **(J)** Time courses for the fidelity of mPFC synaptic inputs onto $LHA_{VGLUT2}$ or $LHA_{VGAT}$ neurons in brain slices, during continuous 1, 5 or 10 Hz optogenetic stimulation for 10 minutes. For 5 Hz: Two-Way Repeated Measures ANOVA, interaction and both main effects $p>0.14$). For 10 Hz: Two-Way Repeated Measures ANOVA, main effect cell type $F(1,6)=132$, $p<0.0001$). *$p<0.05$, **$p<0.01$, ***$p<0.001$.

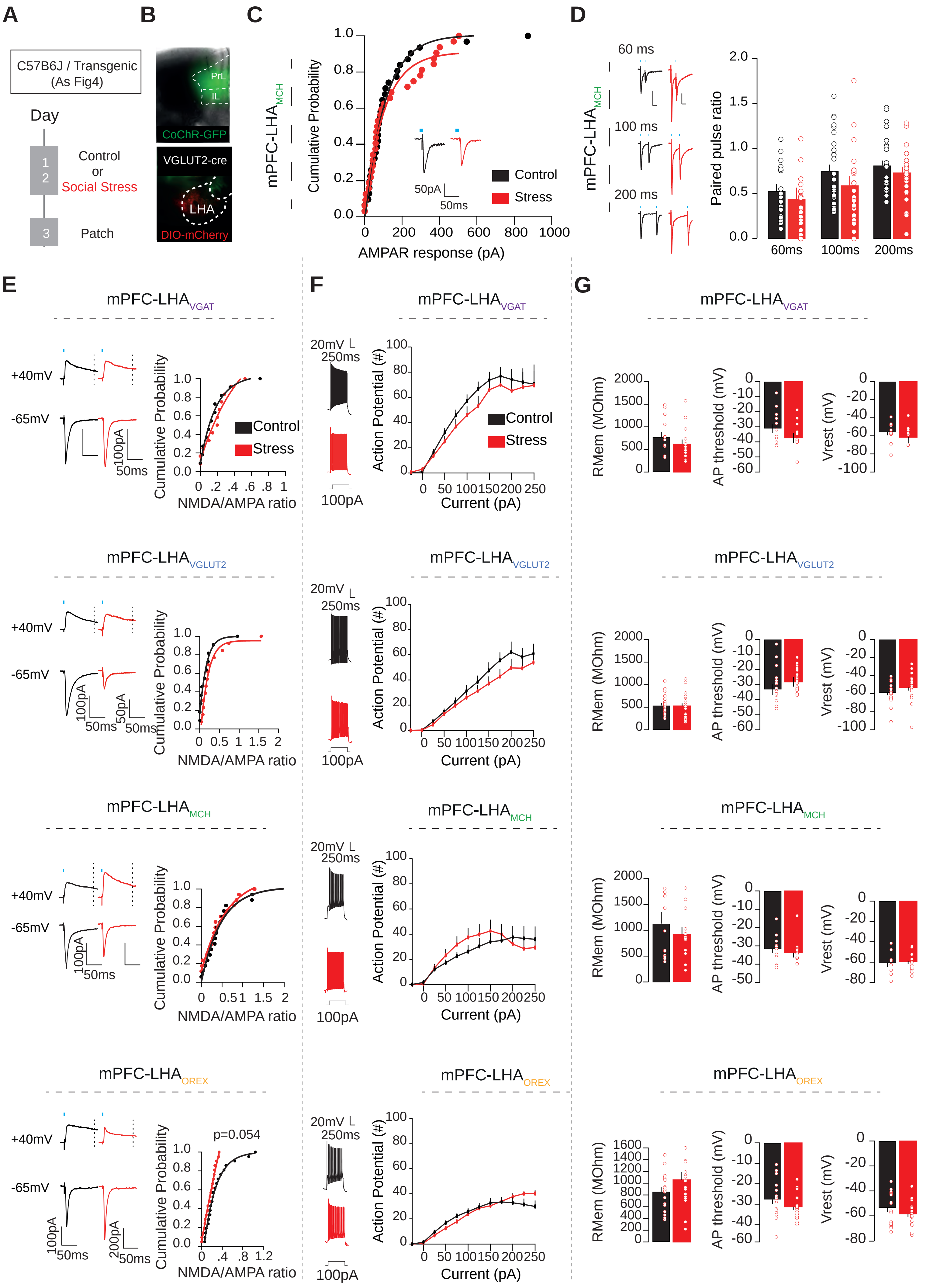


Supplementary Figure 5

**Supplementary Figure 5. Stress does not affect postsynaptic glutamate receptor composition onto mPFC-innervated LHA neuronal subsets or their intrinsic membrane properties. (A)** Schematic and timeline for patch clamp electrophysiology. (**B)** Representative histology image for opsin in mPFC (green) and LHA population identity (red), in a VGLUT2-Cre mouse. Scale bar 200µm**. (C)** Cumulative probability plot for opto-evoked AMPAR amplitudes at mPFC-$LHA_{MCH}$ population (n control=31, 10 mice, n stress=32, 11 mice; Mann-Whitney test, U=486, p=0.894). **(D)** Example traces (left) and Bar graphs (right) for PPR at mPFC-$LHA_{MCH}$ synapses (n control=28, 10 mice, n stress=25, 10 mice; Two-Way RM ANOVA, Main effect of group, F(1,51)=0.001, p=0.969). **(E)** Example traces (-65 mV AMPAR and +40 mV NMDAR at 100 ms vertical dashed line). For mPFC-$LHA_{VGAT}$ synapses: n control=11, 7 mice, n stress=12, 9 mice; Mann-Whitney test, U=55, p=0.5249). For mPFC-$LHA_{VGLUT2}$ synapses: n control=11, 7 mice, n stress=13, 9 mice; Mann-Whitney test, U=49, p=0.2066). For mPFC-$LHA_{MCH}$ synapses: n control=17, 7 mice, n stress= 17, 6 mice; Mann-Whitney test, U=129, p=0.6097). For mPFC-$LHA_{OREX}$ synapses (n control=21, 9 mice, n stress=27, 7 mice; Mann-Whitney test, U=191, p=0.0548). **(F)** Example traces and Current-Spike plots. For mPFC-innervated $LHA_{VGAT}$ neurons (n control=13, 7 mice, n stress=13, 8 mice; Two-way RM ANOVA, Main effect of group, F(1,24)=0.505, p=0.484). For mPFC-innervated $LHA_{VGLUT2}$ neurons: n control=20, 8 mice, n stress=20, 7 mice; Two-way RM ANOVA, Main effect of group, F(1,38)=1.207, p=0.279). For mPFC-innervated $LHA_{MCH}$ neurons: (n control=12, 7 mice, n stress=12, 5 mice; Two-way RM ANOVA, Main effect of group, F(1,22)=0.140, p=0.712). For mPFC-innervated $LHA_{OREX}$ neurons (n control=18, 5 mice, n stress=17, 4 mice; Two-way RM ANOVA, Main effect of group, F(1,33)=0.047, p=0.830). **(G)** Bar graphs for intrinsic membrane properties: membrane resistance (Rmem), action potential threshold and resting membrane potential in control (black) and stress (red) conditions for the distinct LHA cell types. No differences were found. Statistical parameters and group sizes described in the Supplementary statistics table and source files.

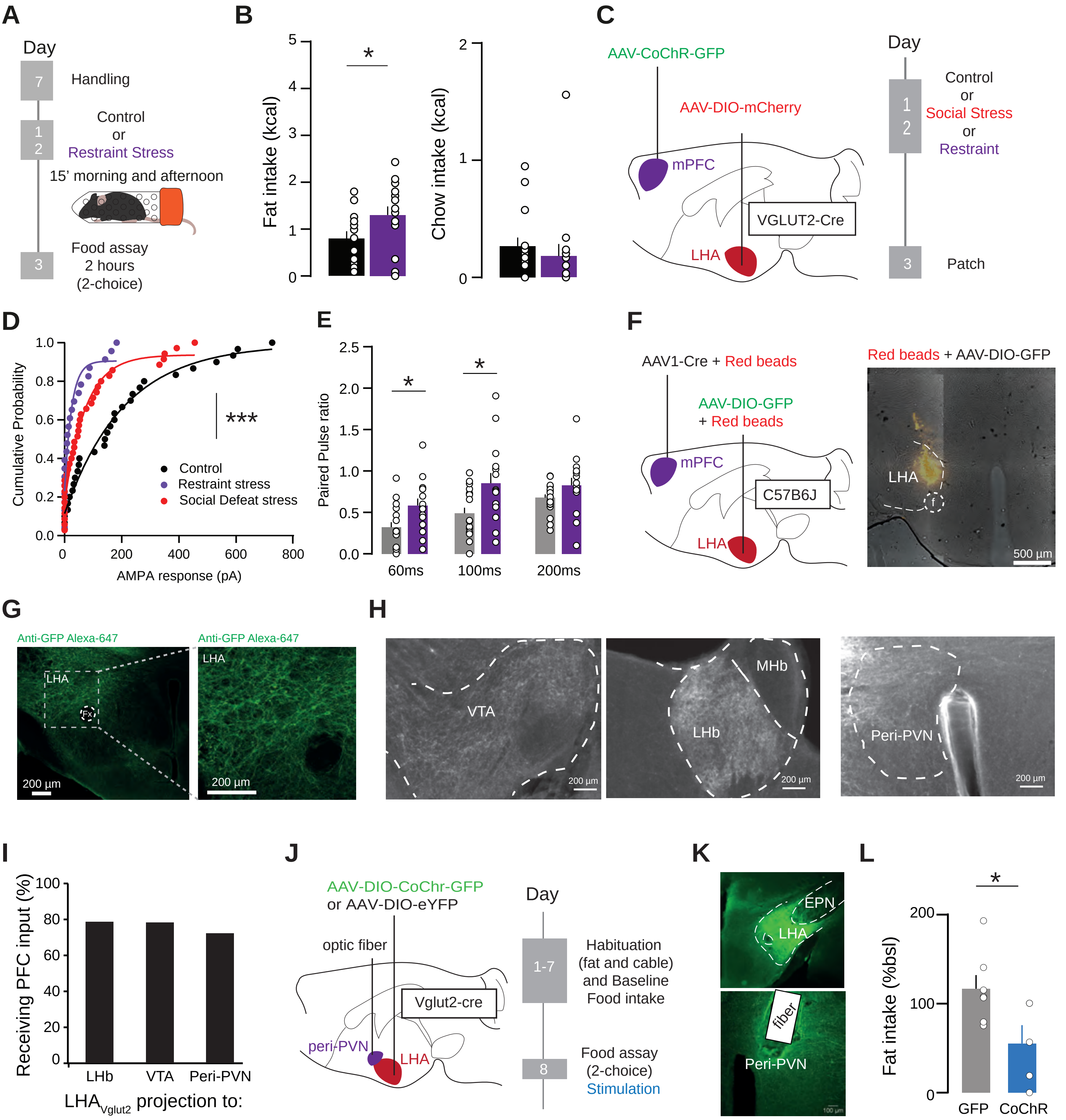


Supplementary Figure 6

**Supplementary Figure 6. Effects of restraint stress and mapping of downstream targets of mPFC-innervated LHA neurons. (A)** Timeline for experiment assessing effect of restraint stress on food intake. **(B)** Quantification of fat (left) and chow (right) intake after restraint stress (purple) or control (black) conditions. Fat: n control=15 mice, n restraint=15 mice, unpaired t test, t(28)= 2.085, p=0.0463. Chow: control n=15, restraint n=15, unpaired t test, t(28)= 0.662, p=0.5134). **(C)** Experimental schematic for evaluating effect of restraint stress on mPFC-$LHA_{VGLUT2}$ synapses. **(D)** Cumulative probability plot for AMPAR response amplitudes after restraint (purple), control (black): (n control=30, 10 mice, n stress=35, 11 mice, n restraint=23, 5 mice; Two-way ANOVA, Main effect of group, F(2;84)=15.05, p<0.0001). **(E)** Bar graphs for PPR effects of restraint stress (n control=20, 10 mice, n stress=19, 11 mice, n restraint= 15, 5 mice; Two-way RM ANOVA, Main effect of group, F(1.89;96.81)=73.36, p<0.0001). **(F)** Left: Viral strategy to inject mPFC neurons with anterograde synapse jumping tracer AAV1-Cre in the mPFC. Subsequent injection of an AAV-DIO-GFP (and red beads for marking the injection site) in the LHA, allowing recombination in mPFC-innervated LHA cells. Right: Image showing stereotactic targeting of the LHA (red beads). **(G)** Recombination of the AAV-DIO-GFP virus in the LHA. **(H)** Image showing fluorescent axons in the VTA, the LHb, and the peri-PVN. Scale bar 200µm. **(I)** Bar graph showing the extent through which $LHA_{VGLUT2}$ neurons with distinct downstream targets received mPFC synaptic input. For $LHA_{VGLUT2}$-LHb: n=15/19 cells connected (79.0%). For $LHA_{VGLUT2}$-VTA: n=51/65 cells connected (78.5%). For $LHA_{VGLUT2}$-Peri-PVN: n=34/47 cells connected (72.3%). **(J)** Schematic showing viral approach (left) and behavioral paradigm (right) to evaluate the effect of optogenetic stimulation of the $LHA_{VGLUT2}$→peri-PVN pathway on food intake. **(K)** Histological validation for $LHA_{VGLUT2}$→peri-PVN pathway stimulation. CoChR (green) expression in the LHA (top) and fiber optic placement above the peri-PVN area and LHA axons (bottom). **(L)** Bar graph quantification of the decreased fat intake after stimulation of this pathway (n GFP=7 mice, n CoChR=5; Mann-Whitney test, U=4, p = 0.028)**.** *= p<0.05; ***= p<0.001.

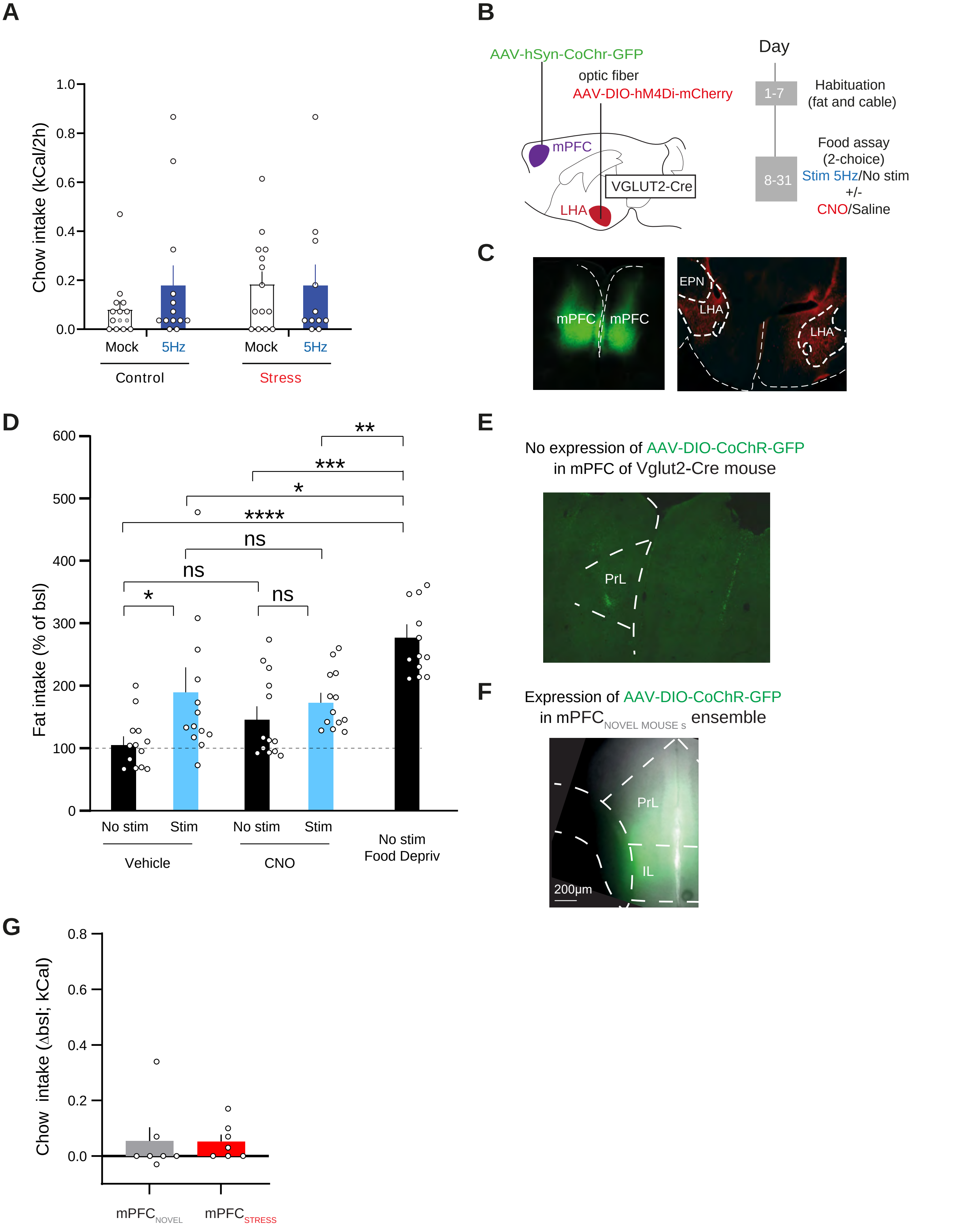


Supplementary Figure 7

**Supplemental Figure 7.** ***In vivo*** **mPFC-LHA optogenetic stimulation and chemogenetic inhibition.** **(A)** Bar graph showing chow intake after stimulation of mPFC-LHA in control and stress conditions (Control-Mock n=14, Control-5Hz n=13, Stress-Mock n=12, Stress-5Hz n=11, Two-way ANOVA, interaction Stimulation-Treatment, $F(1,46)=1.15$, $p=0.29$). **(B)** Schematic for viral strategy (left) and behavioral approach (right) for optogenetic stimulation of mPFC-LHA in combination with chemogenetic inhibition of $LHA_{VGLUT2}$ cells. **(C)** Histological validation. Left: targeting CoChR-GFP in the mPFC. Right: targeting hM4Di-mCherry in the LHA. **(D)** Bar graph of Fat intake (% of baseline) in conditions of mPFC-LHA optogenetic stimulation with or without chemogenetic inhibition of $LHA_{VGLUT2}$ neurons. Right: the percentage-wise increase in fat intake in food deprived animals. Vehicle-NoStim n=13, Vehicle-5Hz n=13, CNO-NoStim n=13, CNO-Stim n=13. Two-Way ANOVA, Stress-CNO interaction, $F(1,24)=2.206$, $p=0.15$. Main effect stim, $p=0.0356$. Posthoc contrasts. Saline: No stim vs Stim, $p=0.01$. CNO: No stim vs stim, $p=0.40$. Saline-NoStim vs CNO-NoStim, $p=0.15$. Food deprivation vs CNO-Stim, $p=0.004$. **(E)** Representative image showing that in a VGLUT2-Cre mouse there is no cre-dependent expression of an opsin in the mPFC. Scale bar 200µm. **(F)** Representative image of opsin expression after TRAP recombination after a novel mouse experience in mPFC. Scale bar 200µm. **(G)** Quantification of chow intake over the 2 hours for experimental groups. Optogenetic stimulation of the $mPFC_{SOCIAL\ STRESS}$ ensemble did not differently affect chow intake from stimulation of the $mPFC_{NOVEL\ MOUSE}$ ensemble projection to LHA (n=7). Mann Whitney $U=17.5$, $p=0.3718$. *= $p<0.05$; ***= $p<0.001$.
